# Phosphorylation regulates arginine-rich RNA-binding protein solubility and oligomerization

**DOI:** 10.1101/2021.06.30.450578

**Authors:** Sean R. Kundinger, Eric B. Dammer, Luming Yin, Cheyenne Hurst, Lingyan Ping, Duc M. Duong, Nicholas T. Seyfried

## Abstract

Post-translational modifications (PTMs) within splicing factor RNA-binding proteins (RBPs), such as phosphorylation, regulate several critical steps in RNA metabolism including spliceosome assembly, alternative splicing and mRNA export. Notably, the arginine-/serine-rich (RS) domains in SR proteins are densely modified by phosphorylation compared with the remainder of the proteome. Previously, we showed that dephosphorylation of SRSF2 regulated increased interactions with similar arginine-rich RBPs U1-70K and LUC7L3. In this work, we dephosphorylated nuclear extracts using phosphatase *in vitro* and analyzed equal amounts of detergent-soluble and -insoluble fractions by mass spectrometry-based proteomics. Correlation network analysis resolved 27 distinct modules of differentially soluble nucleoplasm proteins. We found classes of arginine-rich RBPs that decrease in solubility following dephosphorylation and enrich to the insoluble pelleted fraction, including the SR protein family and the SR-like LUC7L RBP family. Importantly, increased insolubility was not observed across broad classes of RBPs. Phosphorylation regulated SRSF2 structure, as dephosphorylated SRSF2 formed high molecular weight oligomeric species *in vitro*. Reciprocally, phosphorylation of SRSF2 by serine-/arginine protein kinase 2 (SRPK2) *in vitro* prevented high molecular weight SRSF2 species formation. Furthermore, we pharmacologically inhibited SRPKs in mammalian cells and observed increased cytoplasmic granules as well as the formation of cytoplasmic SRSF2 tubular structures that associate with microtubules by immunocytochemical staining. Collectively, these findings demonstrate that phosphorylation may be a critical modification that prevents arginine-rich RBP insolubility and oligomerization.

## INTRODUCTION

RNA-binding proteins (RBPs) cooperatively engage both RNA and protein (1). RBPs frequently contain an RNA-binding domain, typically K-homology (KH) or RNA recognition motif (RRM) domains, that allow the RBP to achieve sequence-specific binding to target RNA molecules (2). Unbiased RNA interactome studies have identified many RBPs containing low complexity (LC) domains that participate in both RNA and protein interactions (2-5). LC domains are typically composed of a select few residues out of the entire amino acid code, giving rise to protein domains that are intrinsically disordered (6). However, LC RBPs exist in a dynamic continuum of native states that range from soluble monomers to liquid-liquid phase separated (LLPS) granules to insoluble fibrils (7) *in vitro* and *in vivo* (8). These assembly states are believed to be influenced in large part by RNA molecules (9,10) and post-translational modifications (PTMs) (11,12). Although we are beginning to decipher a “molecular grammar” regulating LLPS (13), the conditions that give rise to irreversible aggregation are incompletely known.

Recently it has been discovered that the progression of several neurodegenerative diseases is promoted by the aggregation of RBPs (14-21). Interestingly, LC domains are necessary for RBP LLPS and fibrillization (16,19,22,23), processes found to be regulated by PTM. LC RBPs are commonly modified by reversible PTM in the physiological milieu (13,24), yet in neurodegenerative disease phosphorylation PTMs increasingly occupy RBPs such as TDP-43 (25-27). It remains unclear whether phosphorylation is a trigger, or rather a consequence, of pathogenic RBP aggregation.

A major gap in our understanding of arginine-rich proteins is our inability to accurately measure absolute, site-specific phosphorylation levels. Recently our group used middle-down proteomic approaches to demonstrate arginine-rich RBPs have high steady-state levels of PTMs, particularly phosphorylation (28). One such group of arginine-rich RBPs with high levels of phosphorylation is the serine-/arginine-rich (SR) splicing factor family of RBPs (29). This twelve member RBP family is known to contain at least one RRM RNA-binding domain (30) at the N-terminus and a C-terminal arginine-/serine-rich (RS) domain distinguished by an expanded tract of RS dipeptide motifs, a phosphomotif conserved from yeast to man (3). The most extensively studied regulator of SR protein function is phosphorylation, primarily catalyzed by nuclear cdc2-like kinases (CLKs) (31) and cytoplasmic SR protein kinases (SRPKs) (32-34). Phosphorylation regulates nearly every facet of SR protein function, including splicing (35), coupling to sites of active transcription (36,37), subcellular localization (38-40), nuclear speckle compartmentalization (31,32,41) and binding partner selection and affinity (30,39,42,43). Importantly, it is not fully understood whether excessive, or rather, insufficient phosphorylation alters the stability of SR proteins.

Our group (28) and others (40,44-46) suggest SR proteins may increasingly bind together and aggregate when insufficiently phosphorylated. Importantly, SR proteins and proteins that harbor homologous domains can aggregate under native conditions (47). Collectively, these data support a hypothesis that dephosphorylation would result in SR proteins becoming insoluble, as well as those RBPs with SR-like LC domains.

Here, we sought to understand the role of phosphorylation in regulating RBP solubility. We enriched for RBPs by biochemical fractionation from mammalian cell lines and incubated with calf intestinal alkaline phosphatase (CIP), which catalyzes the removal of phosphate PTMs from proteins (48). We conducted liquid-chromatography coupled with tandem mass spectrometry (LC-MS/MS) on detergent-soluble and -insoluble pellet fractions of dephosphorylated and mock-treated nucleoplasm extracts and used a network based approach to identify groups of RBPs that exhibited similar solubility changes that were regulated by phosphorylation. Importantly, we found that SRSF2 and related SR proteins co-aggregated to the insoluble fraction, while other nuclear RBPs such as TDP-43 did not. Moreover, we found that phosphorylation regulates SRSF2 assembly states *in vitro*. Finally, we show that SRPK inhibition in cells results in an increase in the number of cells harboring cytoplasmic SRSF2 granules as well as filamentous-like structures that co-localize with microtubules. Collectively, this work reinforces phosphorylation as an important regulator of SR protein solubility and structure and suggests that phosphorylation may be a preventative cellular mechanism against arginine-rich RBP aggregation.

## RESULTS

### Phosphorylation prevents SRSF2 aggregation

Phosphorylation is a critical regulator of RNA-binding protein (RBP) protein function (45,49), condensation (12,50,51) and overall structure (52,53). The serine-/arginine-rich (SR) splicing factor family of RBPs is observed to be substantially phosphorylated, although a wide range of phosphorylation states are predicted for SR proteins *in vivo* (28). Recently, we reported that dephosphorylation of serine-/arginine-rich splicing factor 2 (SRSF2), a prototypical arginine-rich nuclear speckle phosphoprotein (28,33,54,55), enhanced interactions with fellow arginine-rich RBPs (28), leading us to hypothesize that phosphorylation regulates the solubility of structurally similar arginine-rich RBPs. Here, we use SRSF2 as a paradigm to study the regulation of arginine-rich RBP solubility, structure and morphology by phosphorylation. In SRSF2, the RS domain is highly phosphorylated (28), a region with high probability of intrinsic disorder (**Supplemental Fig.S1a**) (56,57). We incubated lysates containing recombinant SRSF2-myc, a known phosphoprotein, with calf intestinal alkaline phosphatase (CIP) which corresponded to a faster migrating SRSF2 band (**Supplemental Fig.S1b**) by SDS-PAGE, suggesting substantial dephosphorylation of SRSF2 (58,59). To further validate dephosphorylation of SRSF2, we immunoblotted with an antibody raised against the C-terminus of SRSF2 that preferentially labels hypophosphorylated SRSF2 (*hypo*SRSF2) (60-62) (**Supplemental Fig.1c**). We once again observed increased migration of SRSF2. Furthermore, we saw an increase in *hypo*SRSF2 labeling, demonstrating that SRSF2 is indeed dephosphorylated.

We then asked whether phosphorylation regulates the solubility of SRSF2. Detergent-soluble (S) and -insoluble pelleted (P) fractions were isolated following mock (-CIP) or phosphatase (+CIP) treatment (**Fig.1a**). We resolved equal amounts of total (T), soluble (S) and pelleted (P) fractions by SDS-PAGE and immunoblotted for SRSF2-myc (**Fig.1b**). While phosphorylated SRSF2-myc was primarily soluble (68% of total) in the mock condition, dephosphorylated SRSF2-myc significantly decreased in solubility, enriching to the detergent-insoluble pellet fraction (89% of total, **Fig.1c**). This suggests similar arginine-rich RBPs or groups of RBPs may experience altered solubility following dephosphorylation as well. We next sought to globally identify and quantify RBPs that aggregate following dephosphorylation.

**Figure 1.**
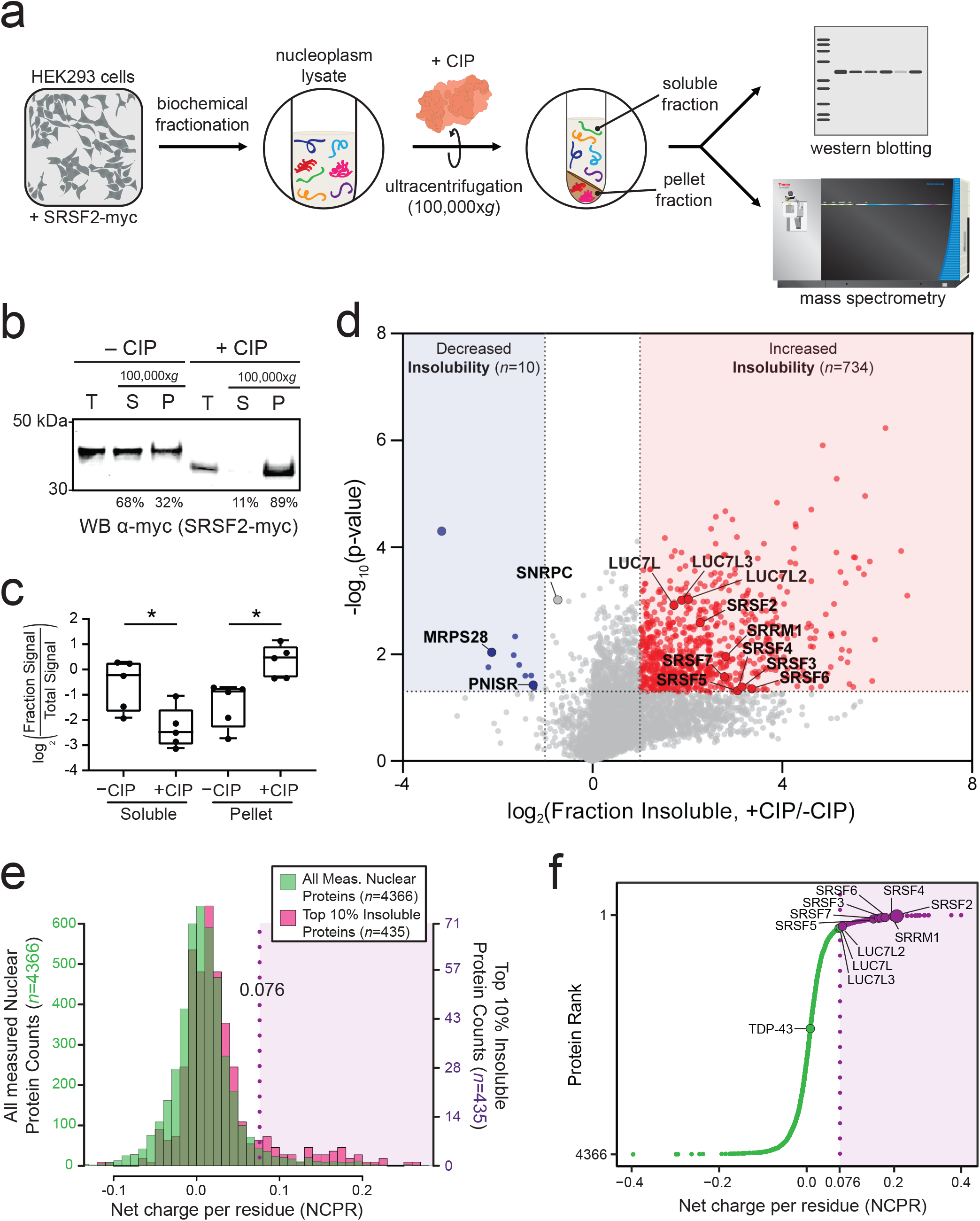
Phosphorylation regulates arginine-rich RNA-binding protein solubility. **(a)** Sample preparation and proteomic workflow. Nucleoplasm extracts of HEK293 cells expressing recombinant SRSF2-myc protein were incubated with either calf intestinal phosphatase (+CIP) or distilled water (-CIP) at 37°C for 1 hour. Following this, samples were ultracentrifuged at 100,000x*g* for 1 hour. The soluble and insoluble pellet fractions were desalted and run by either western blot or liquid chromatography coupled with tandem mass spectrometry. **(b)** Pre-spin input (total, *T*), supernatant (soluble, *S*) and insoluble pellets (*P*) were run by SDS-PAGE and western blotted for myc. The average percent soluble (*sol*.*/(sol. + insol*.)] and insoluble (in*sol*.*/(sol. + insol*.)] values were calculated for five biological replicates and displayed below the representative western blot. **(c)** Band densitometry of soluble and pellet fraction log2-transformed SRSF2-myc band intensities normalized to the total signal in -CIP and +CIP conditions (5 biological replicates; Soluble p value = 0.0123; Pellet p value = 0.0372; two-tailed paired t-test). **(d)** Differential abundance of proteins in the soluble fractions. Fold-change, displayed on the x-axis, was the log2 value for fraction of signal that was insoluble [insoluble/(insoluble+soluble)] for the pairwise comparison +CIP/-CIP. The t-statistic (-log10(p-Value)) was calculated for all proteins and displayed on the y-axis. Insoluble-enriched proteins were highlighted in *red* (log2(fold change) ≥ 1, p value < 0.05) and proteins depleted from the insoluble fractions upon dephosphorylation were highlighted in *blue* (log2(fold change) ≤ -1, p-Value < 0.05) squares, respectively. **(e)** Histogram plot of the net charge per residue (NCPR) of all measured nuclear proteins (*n*=4366; *green*) overlaid with the top 10% most insoluble proteins (*n*=435; *magenta*). The left y-axis corresponds to protein bin counts of all measured nuclear proteins (*green*) whereas the right y-axis corresponds to the protein bin counts of the top 10% insoluble proteins (*magenta*). Gaussian curves were fit to the histogram plots, with two curves fit to the top 10% insoluble proteins. The NCPR range right of the right-most gaussian curve mid-point (NCPR=0.076) was shaded *purple*. **(f)** An S-graph ranking each protein by the NCPR value. SR proteins are highlighted, ranking among the highest NCPR value proteins in the proteome. The region of proteins with NCPR values greater than 0.076 was shaded *purple*, while proteins with NCPR<0.076 were colored *green*.

### Proteomics reveals RBPs that aggregate following dephosphorylation

The soluble and pellet samples were analyzed by label-free quantitative proteomics, using liquid chromatography coupled with tandem mass spectrometry (LC-MS/MS) (**Fig.1a, Supplemental Table S1**). Notably, dephosphorylation did not induce global aggregation of the nuclear proteome, as insoluble pellet fraction protein concentrations were unchanged after phosphatase incubation (**Supplemental Table S2**). Following database search and removal of proteins with >50% missing values, we identified 4,366 unique proteins (**Supplemental Table S3**).

To discover proteins with the largest changes in solubility following dephosphorylation, we calculated the log2 fold differences of fraction insoluble values between phosphatase and mock treatments and visualized this as a volcano plot (**Fig.1d, Supplemental Tables S4-5**). Proteins were highlighted as enriched/depleted from a fraction if a greater than or equal to a two-fold change in fraction insoluble values was observed with a p value less than 0.05 (two-tailed paired t-test). Relatively few proteins (*n*=10) enriched to the soluble fraction following dephosphorylation. In contrast, many more proteins (*n*=734) experienced increased aggregation following dephosphorylation. Among these were SRSF2 and fellow SR proteins (SRSF3/4/5/6/7), as well as the SR-like SRRM1 and LUC7L RBP family. These observations suggest phosphorylation is an important PTM that regulates the solubility of SRSF2, as well as the solubilities of similar arginine-/serine-rich RBPs.

### Arginine-/lysine-rich RNA-binding proteins with positive net charge preferentially aggregate following dephosphorylation

Recently, the Jedd group generated a synthetic, recombinant arginine-rich low complexity protein equally balanced in positively-charged and negatively-charged residues in a dipeptide repeat pattern, and found that increasingly incorporating positively-charged residues resulted in increased aggregation of the recombinant protein (44). This suggests that increased net positive charge by virtue of increased arginine and lysine content may increase the condensation of low complexity RBPs. Importantly, phosphorylation significantly alters the net charge of a protein, adding a -2 charge with each phosphorylated residue at physiological pH (63). We hypothesized that proteins with highly positive net charge (high densities of arginine/lysine) may be predisposed to aggregate when not sufficiently phosphorylated.

We generated separate histograms of net charge per residue (NCPR) values for the nuclear proteome sequenced (*n*=4,366 proteins) and of those, the top 10% most insoluble proteins (*n*=435 proteins) following dephosphorylation (**Fig.1e**). The average NCPR of the top 10% most insoluble proteins (0.028) was significantly increased from the proteome studied (0.009) (p value = 8.2e-15, Welch Two Sample t-test). The top 10% most insoluble proteins exhibited a bimodal distribution of NCPR, with a shoulder of proteins with a highly positive mean NCPR value (0.076). These data suggest proteins with a high NCPR (> 0.076) are susceptible to increased aggregation following dephosphorylation. We then plotted the distribution of NCPR of the nucleoplasm proteome sequenced and highlighted those proteins with NCPR>0.076 (**Fig.1f**). Interestingly, 11/12 members of the SR protein family surpass this threshold, with SRSF2 being the most positively charged overall. The SR RBP family has an abnormally high positive average net charge, relative to other RBP families (**Supplemental Table S6**). SRSF2, as well as SRSF3/4/5/6/7 all experienced significantly aggregation following dephosphorylation. This suggests that phosphorylation may be an especially important mechanism to regulate the solubility of proteins with high concentrations of arginine and lysine, among which are many nuclear RBPs involved in splicing.

### Systems analysis identifies modules of proteins with solubility impacted by phosphorylation

We hypothesized that if we applied systems biology approaches to the protein abundance data we collected, we could discover groups of structurally-similar proteins that may co-aggregate when not sufficiently phosphorylated. To test this, we performed Weighted Gene Correlation Network Analysis (WGCNA) (64) to group proteins with highly correlated soluble and insoluble fraction abundance patterns. To define functionally divergent protein groups, we plotted a dendrogram that was segregated by hierarchical clustering into modules of related proteins (**Supplemental Fig.S2**). The network reduced our proteome into 27 modules [rank ordered by size, M1 (largest) – M27 (smallest)] each assigned a representative color (**Supplemental Table S7**). Each module was classified by the strength of associations to GO terms linked to discrete and generalizable cellular functions.

To understand which module of proteins experienced the greatest alterations in solubility following dephosphorylation, we performed a one-dimensional hypergeometric Fisher’s exact test (FET) for enrichment within each module of those proteins with the most increased insolubility following CIP-treatment (*n*=734) (**Fig.2a**). Several key modules (M5, M7, M16) were enriched with proteins that aggregated following dephosphorylation. To identify protein drivers behind module solubility changes, eigenprotein values were plotted according to protein fraction and treatment condition (**Fig.2b-c**). Eigenproteins are defined as the first principal component of a module and serve as a representative, weighted module expression profile. As expected, some modules did not experience appreciable solubility changes following dephosphorylation. Among these were the soluble module M1 (protein folding) and insoluble module M9 (mRNA splicing), both unchanged in solubility profile after dephosphorylation (**Fig.2b**). Other modules, however, including M5 (dephosphorylation), M7 (cell cycle phase) and M16 (DNA repair) represented a prominent group of modules comprised of proteins that are soluble when phosphorylated, yet aggregate upon dephosphorylation (**Fig.2c**). The three members of the SR-like LUC7L family were each hub proteins of the M7 module, which experienced a large decrease in solubility. SRSF2 was a member of the M16 module, which interestingly also contained several cytoskeletal components (**Supplemental Table S7**). Given the robust insolubility of individual modules following dephosphorylation, we sought to validate individual proteins that significantly changed in solubility as well.

**Figure 2.**
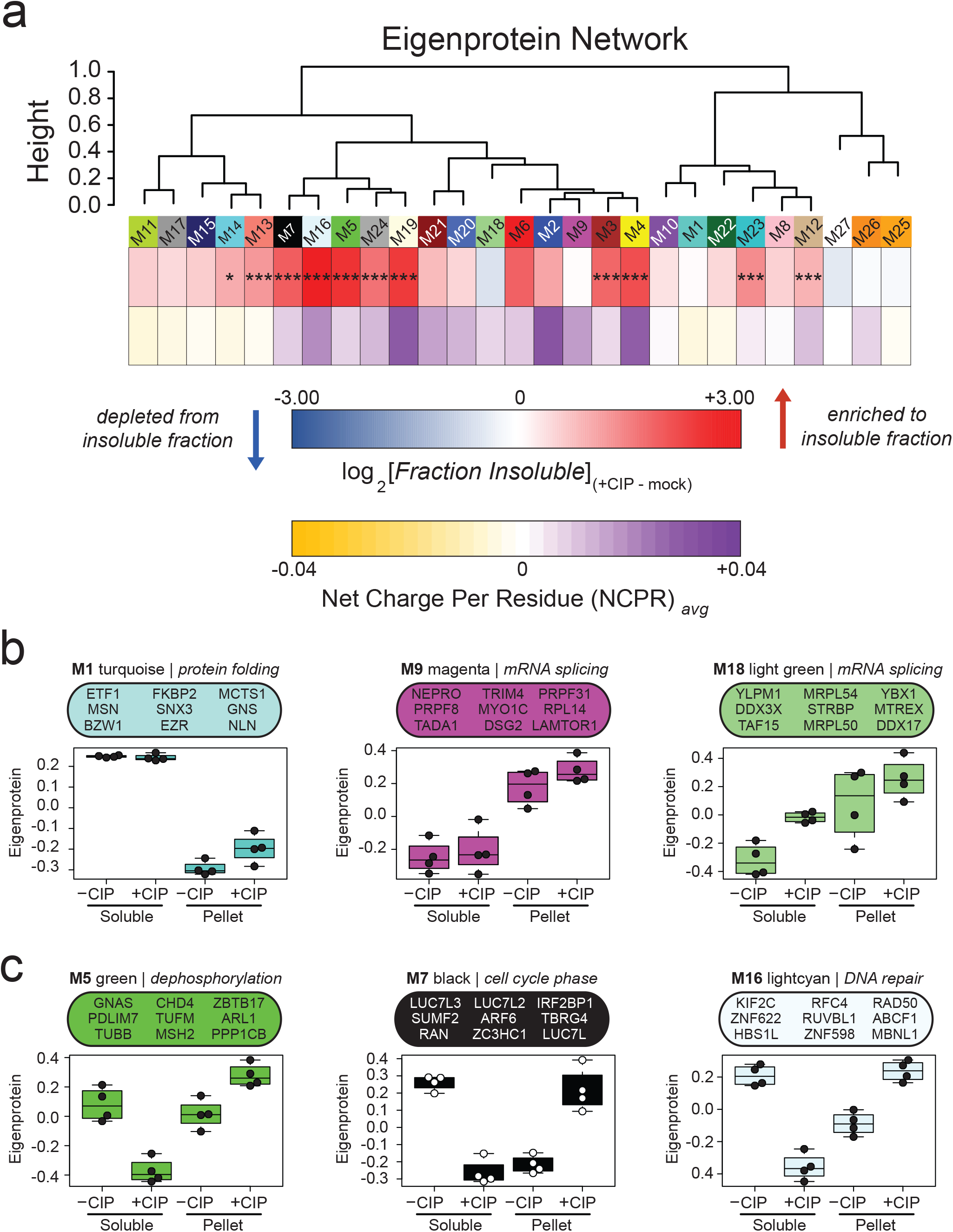
A correlation network approach groups proteins into modules with discrete gene ontologies and solubility patterns in response to dephosphorylation. **(a)** Protein modules were clustered to assess module relatedness based on correlation with abundances in detergent-soluble and -insoluble pellet fractions in -CIP and +CIP conditions. Log2 values for fraction of signal that was insoluble [insoluble/(insoluble+soluble)] for the pairwise comparison +CIP/-CIP displayed as a heatmap for each module and compared with the average net charge per residue (NCPR) of all members of that module, also displayed as a heat map. Significance of enrichment to top 734 insoluble proteins displayed as asterisks, determined by one-dimensional hypergeometric Fisher’s exact test (FET, BH corrected; *p < 0.05, **p < 0.01, ***p < 0.001). Modules with positive average NCPR are colored *purple* while those with negative average NCPR are colored *yellow*. **(b-c)** Eigenprotein abundance box and whisker plots of selected modules. The farthest data points, up to 1.5 times the interquartile range away from box edges, define the extent of whiskers (error bars). **(b)** Protein modules that are unchanged in solubility phosphorylated or dephosphorylated. **(c)** Selected modules with increased insolubility abundances following dephosphorylation.

### Confirmation of hub protein solubility changes following dephosphorylation

To examine the effect of dephosphorylation on the solubility of individual RBPs, we plotted the mass spectrometry protein abundance measurements (**Fig.3a**) and compared with immunoblot signals of select endogenous proteins from total, soluble and pelleted fractions (**Fig.3b**). The classical SR proteins SRSF1 and SRSF2 were depleted from the soluble fraction which was verified by western blot. LUC7L and LUC7L3, both hubs of the M7 ‘cell cycle phase’ module, were significantly depleted from soluble fractions and enriched to insoluble pellet fractions. Interestingly, the U1 small nuclear ribonucleoprotein C (SNRNPC), a member of the U1 snRNP subunit of the spliceosome, experienced increased solubility following dephosphorylation. Other nuclear RBPs in HEK293 nucleoplasm lysates including TDP-43 (M5), hnRNPAB (M21) and ZC3H18 (M9), however, showed no significant solubility changes following dephosphorylation. Mass spectrometry examination of soluble and pellet fractions revealed global RBP solubility changes in response to phosphatase co-incubation, revealing RBPs susceptible to destabilization following dephosphorylation.

**Figure 3.**
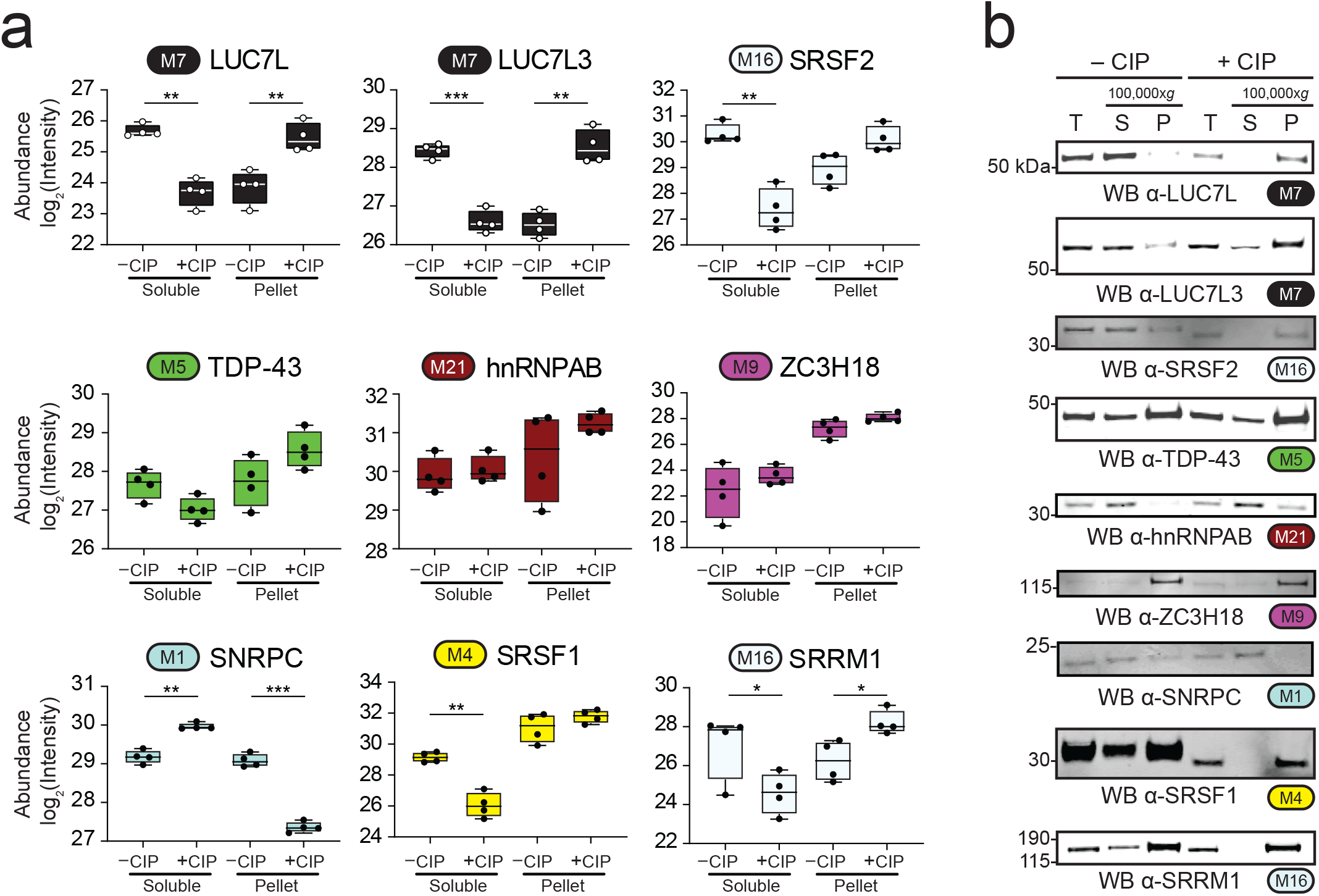
RNA-binding proteins have variable abundance patterns in soluble and pellet fractions following dephosphorylation. **(a)** Box and whisker plots of mass spectrometry abundance measurements (*n*=4) of RNA-binding proteins in soluble and pellet fractions in -CIP and +CIP conditions. (two-tailed paired t-test; *p < 0.05, **p < 0.01, ***p < 0.001). Module number and color are indicated next to each gene symbol. Whiskers range from min to max values. **(b)** Western blot validation of solubility changes of well-described RNA-binding proteins. Modules, colored accordingly, are paired with the protein name.

### Phosphorylation decreases NCPR and regulates the oligomerization of arginine-rich SRSF2

With no consideration of post-translational modifications (PTMs), SRSF2 is among the most positively charged proteins in the entire proteome (**Supplemental Table S8**). In a theoretical exercise of how phosphorylation changes the net charge of the SRSF2, we calculated the average SRSF2 NCPR **(Fig.4a)** and local charge density in a 21 residue sliding window range **(Fig.4b)** assuming three separate states: no phosphorylation (*hypo*SRSF2), MS-observed phosphorylation (pSRSF2†) (28), and full phosphorylation (*hyper*pSRSF2) (**Supplemental Table S9**). Although substantially positively-charged within the RS domain, SRSF2 adopts increasing negative charge with increasing phosphorylation, such that the RS domain becomes net negatively-charged at full phosphorylation occupancy. As we observed increased SRSF2 aggregation upon dephosphorylation, we asked whether phosphorylation could similarly regulate the oligomerization of recombinant SRSF2.

Given the aggregation we observed for SRSF2, we hypothesize that a critical structural change occurs as a result of dephosphorylation. To test this, we analyzed phosphorylated and dephosphorylated lysates containing SRSF2-myc by both denaturing SDS-PAGE and non-denaturing Blue native PAGE followed by western blotting for the myc-tag of recombinant SRSF2 (**Fig.4c-d**). In contrast to SDS-PAGE, which separates proteins under denaturing conditions, native PAGE resolves native protein masses in high molecular weight oligomeric states as well as protein complexes formed by physiological protein-protein interactions (65). The mock-treated SRSF2-myc sample had a recognizable monomer species band at ∼ 37 kDa, whereas dephosphorylated SRSF2 exhibited dimer and tetramer species formation, as well as high molecular weight oligomeric species (**Fig.4d**). These data suggest that dephosphorylation leads to higher molecular weight oligomeric SRSF2 structures.

To reciprocally demonstrate how SRSF2 structure changes with phosphorylation, we performed an *in vitro* kinase reaction using the SR protein kinase 2 (SRPK2) (**Fig.4e-f**) (66). Both SRPK2 and SRSF2 were expressed and purified in *E. coli*, an organism with a low abundance of phosphoproteins and only three known Ser/Thr-directed kinases (67,68). Despite not observing an overt change in molecular mass via SDS-PAGE following phosphorylation (**Supplemental Fig.S3a**), the increased migration of SRSF2 by isoelectric focusing (IEF) (**Supplemental Fig.S3b**) is consistent with reduced net charge and increased phosphorylation. We then separated these reaction products by non-denaturing native PAGE and found that unphosphorylated SRSF2 formed high molecular weight oligomers, with a prominent high molecular weight band at ∼720 kDa (**Fig.4f**). In contrast, SRPK2-phosphorylated SRSF2 existed primarily as monomeric species. These data demonstrate that phosphorylation regulates SRSF2 homotypic interactions and prevents high molecular weight oligomeric species formation.

**Figure 4.**
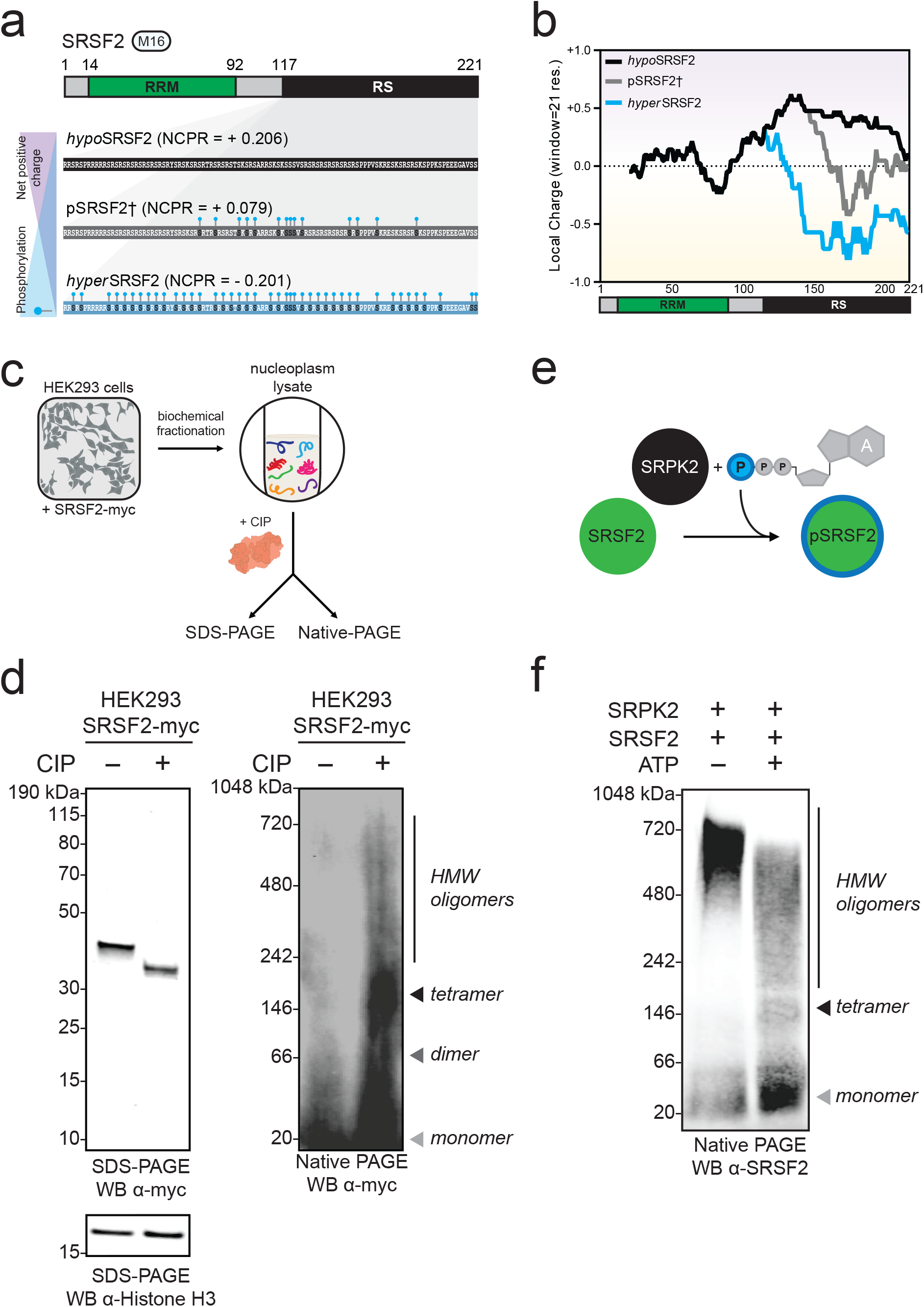
SRSF2 net charge and insolubility, respectively, increases substantially with dephosphorylation. **(a)** SRSF2 net charge per residue (NCPR) calculated according to phosphorylation state, i.e., no phosphorylation (*hypo*SRSF2), phosphorylation sites previously observed by middle-down mass spectrometry (pSRSF2†, Kundinger & Bishof *et al*., 2020) or full phosphorylation (*hyper*pSRSF2). Phosphorylation sites represented by turquoise ball and stick. **(b)** Line plots of average charge density (window = 21 residues) from C-terminus to N-terminus of SRSF2 that is either non-phosphorylated (*black*), observed phosphorylation by MS (*grey*, Kundinger & Bishof *et al*., 2020) or fully phosphorylated (*turquoise*) in the RS domain. The SRSF2 protein map is included below the line plot, with the RNA-recognition motif (RRM) domain (*green box*) and RS domain (*black box*) annotated. **(c)** Nucleoplasm extracts of HEK293 cells expressing recombinant SRSF2-myc protein were incubated with either calf intestinal alkaline phosphatase (+CIP) or distilled water (-CIP) at 37°C for 1 hour. Following this, samples were split and run by both denaturing and non-denaturing native PAGE and western blotted for myc (*n*=3). **(d)** By denaturing SDS-PAGE (*left*), CIP-treated SRSF2-myc has increased electrophoretic mobility. Equal loading is demonstrated by Histone H3 labeling. Immunoblotting for SRSF2-myc after non-denaturing Blue native PAGE (*right*) identifies various dephosphorylated SRSF2 species not observed in mock-treated samples, including monomer (∼37 kDa, *light grey triangle*), dimer (∼74 kDa, *dark grey triangle*), tetramer (∼148 kDa, *black triangle*) and high molecular weight (HMW) species. **(e)** Diagram of *in vitro* kinase reaction using kinase SRPK2 (*black*) and substrate SRSF2 (*green*). Both SRPK2 and SRSF2 were expressed and purified from *E. coli* and were mixed in the presence (+) or lack thereof (-) ATP and incubated at 30°C for 30 min. Phosphorylated SRSF2 was represented by a turquoise ring. **(f)** The *in vitro* reaction was separated by non-denaturing native PAGE which was transferred and western blotted for SRSF2. Monomeric (*grey triangle*), tetrameric (*black triangle*) and high molecular weight (HMW) oligomeric (*line*) species were unequally observed in the – and + ATP conditions by non-denaturing native PAGE.

### SRPK inhibitor SRPIN340 decreases SR protein phosphorylation and increases SRSF2 granule and tubule formation

Having established that SRSF2 aggregates and forms high molecular weight oligomers upon dephosphorylation *in vitro*, we attempted to inhibit SR protein phosphorylation in cells. We incubated HEK293 cells in media containing the compound SRPIN340, a cell-permeable, ATP-competitive selective SRPK inhibitor (69). To validate SRPIN340 we analyzed cell lysates by western blot using mAb104 (70), an antibody that labels phosphoepitopes on numerous SR proteins (71). We also co-labeled with an antibody that recognizes phosphorylated SRSF2 (pSRSF2) epitopes called SC35 (54), a synonym for the SRSF2 gene product protein (**Fig.5a**). Quantification of mAb104 immunoblot signal intensities normalized to Histone H3 demonstrate that several SR proteins, particularly SRSF2, exhibit decreased phosphorylation with increasing concentrations of SRPIN340 (**Fig.5b**). Independently, we compared SC35 labeling of pSRSF2 with an antibody anti-phosphoTDP-43 (pTDP-43). We found that while pTDP-43 did not quantitatively change with SRPIN340 treatment, pSRSF2 signal was significantly decreased at 50 µM SRPIN340 (**Fig.5c**). These results demonstrate that inhibition of SRPKs using SRPIN340 successfully reduces pSRSF2 levels, as well as overall SR protein phosphorylation.

**Figure 5.**
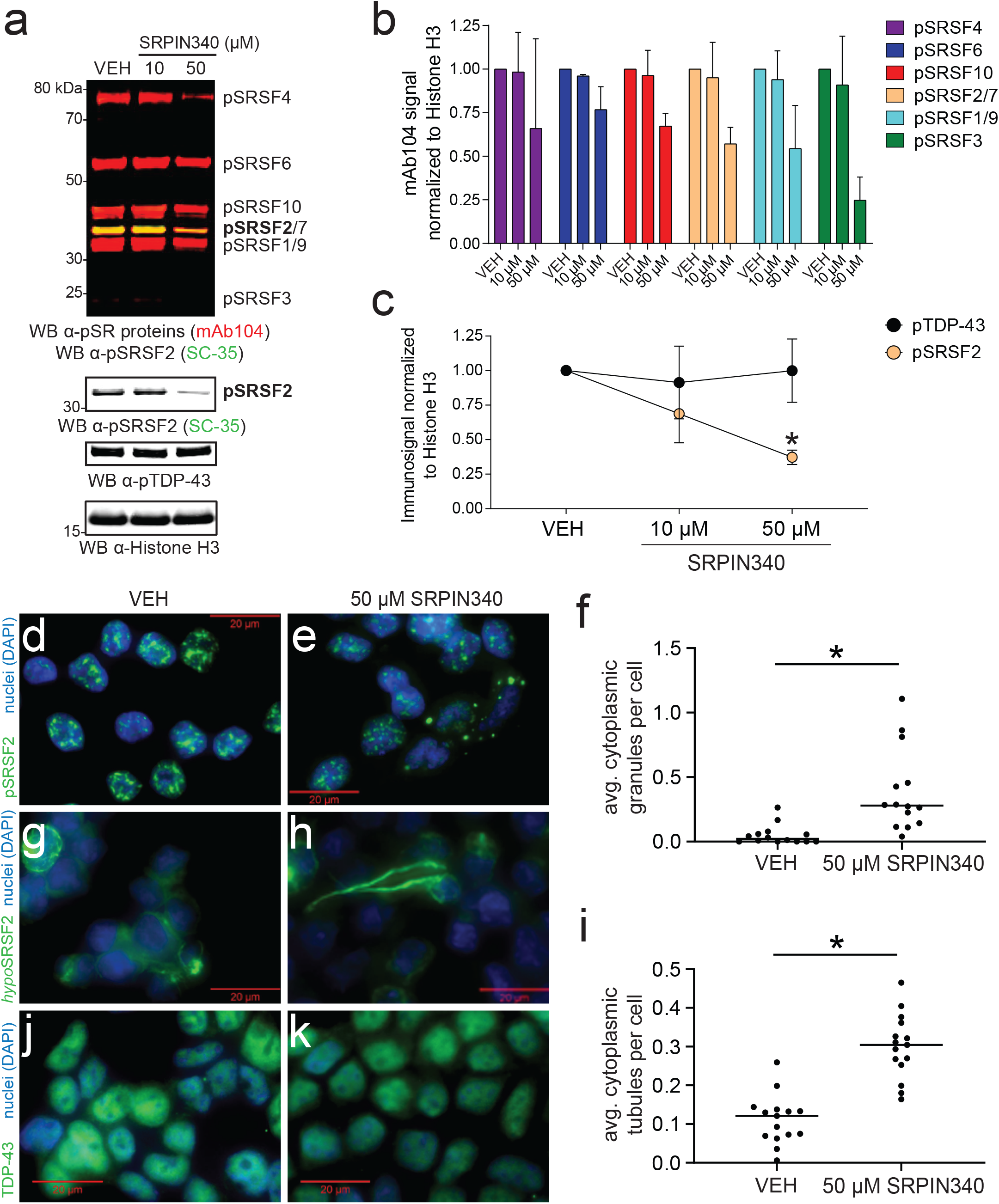
Inhibiting SRPKs decreases SR protein phosphorylation and increases cells harboring cytoplasmic SRSF2 granule and tubule structures in HEK293 cells. **(a)** HEK293 cells were incubated with either DMSO (vehicle, VEH) or increasing concentrations of SRPK inhibitor SRPIN340 for a length of 12 hours. Immediately following treatment, cells were harvested, run by SDS-PAGE and Western blotted for phosphoSR (pSR) proteins SRSF4 (*purple*), SRSF6 (*blue*), SRSF10 (*red*), SRSF2/SRSF7 (*gold*), SRSF1/9 (*turquoise*) and SRSF3 (*green*) (mAb104 antibody, *red signal*). These membranes were also co-labeled using the SC35 antibody (*green signal*) that independently recognizes pSRSF2 (*gold*). The putative pSRSF2 band is indeed labeled by both mAb104 (*red signal*) and SC35 (*green signal*), such that the 35 kDa band appears a *yellow* color, bolstering confidence that pSRSF2 is indeed that protein. A p**-**TDP-43 antibody and a Histone H3 antibody were used to confirm specificity of SRPIN340 and equal protein loading, respectively. **(b)** mAb104 pSR protein band intensities for SRSF4 (*purple*), SRSF6 (*blue*), SRSF10 (*red*), SRSF2/SRSF7 (*gold*), SRSF1/9 (*turquoise*) and SRSF3 (*green*) of SRPIN340 treatments (10 µM, 50 µM) were quantified and normalized to the DMSO condition (artificially set to value=1). Error bars indicate maximum and minimum ranges of band values. **(c)** Band intensities of pTDP-43 (*black*) and pSRSF2 (*gold*) were quantified and normalized to Histone H3 labeling. The DMSO condition was artificially normalized to equal 1.00 and compared with normalized pTDP-43 and pSRSF2 values in SRPIN340 conditions. The pSRSF2 signal was significantly decreased compared with pTDP-43 at 50 µM SRPIN340 concentration (multiple t tests, *p value=0.0099). **(d-e)** Immunocytochemical (ICC) staining of HEK293 cells for phosphoSRSF2 (pSRSF2)-positive nuclear speckles (*green*) and DAPI+ nuclei (*blue*) in vehicle-treated cells **(d)** and 50 µM SRPIN340-treated cells **(e). (f)** The number of cytoplasmic granules observed were divided by the number of cells counted and averaged for 14 independent images in three independent replicates (*p value = 0.0191, two-tailed paired t-test). A minimum of 120 cells were counted in each condition per replicate. The error bars represent the range of standard deviation. **(g-h)** ICC staining of HEK293 cells for hypophosphorylated SRSF2 (hypoSRSF2) (*green*) and DAPI+ nuclei (*blue*) in vehicle-treated cells **(g)** and 50 µM SRPIN340-treated cells **(h). (i)** The fraction of cells harboring cytoplasmic SRSF2 tubule structures was quantified in four biological replicates and compared (*p value = 0.0178, two-tailed paired t-test). **(j)** Vehicle-and **(k**) 50 µM SRPIN340-treated HEK293 cells were stained by ICC for TDP-43 (*green*) and DAPI-stained.

As we observed high molecular weight species of dephosphorylated SRSF2 *in vitro*, we hypothesized that inhibiting SR protein kinases within cells would induce an increase in cytoplasmic granules and/or the formation of fibril-like species, hallmarks of various RBP proteinopathies (72). Immunocytochemical (ICC) staining using the nuclear speckle antibody SC35 demonstrated canonical nuclear speckle morphology of pSRSF2 in vehicle-treated cells (**Fig.5d**). In cells incubated with SRPIN340, we observed unusual cytoplasmic granules (**Fig.5e**). We performed quantified these and observed a significant increase in the average number of cytoplasmic granules per cell in the SRPIN340 condition (**Fig.5f**). We next labeled cells using the *hypo*SRSF2 antibody, observing dephosphorylated SRSF2 dispersed largely throughout the cytoplasm and infrequently observed as small tubules extending from the cell body outwards (**Fig.5g**). In cells incubated with SRPIN340, however, we observed a significant increase in cells harboring cytoplasmic SRSF2 tubule structures, which were substantially larger in size (**Fig.5h-i**). As expected, granule condensation and formation of tubule-like filaments was not observed for TDP-43 which showed a consistent pattern of nuclear distribution in both conditions **(Fig.5j-k**).

### SRSF2 interacts with microtubule subunit proteins α- and β-tubulin

Given the unusual tubular morphology we observed of hypophosphorylated SRSF2 in cells and enhanced aggregation of SRSF2 following dephosphorylation *in vitro*, we further examined constituents of M16, of which SRSF2 is a member **(Fig.6a)**. Interestingly, several proteins within this module are components of the cytoskeleton, including microtubule subunit protein β-tubulin 8 (TUBB8). To answer whether SRSF2 interacts with microtubules, we immunopurified recombinant SRSF2 and immunoblotted for TUBB8 and TUBA1A. We indeed found that SRSF2 interacts with TUBB8 and TUBA1A (**Fig.6b**). Furthermore, ICC staining of SRSF2 with either TUBA1A or TUBB8 demonstrates that dephosphorylated SRSF2 preferentially associates with microtubule structures (**Fig.6c-d**). Thus, network-based proteomics revealed an association between arginine-rich RBPs like SRSF2 and microtubules, confirmed by both biochemical and cell imaging approaches.

**Figure 6.**
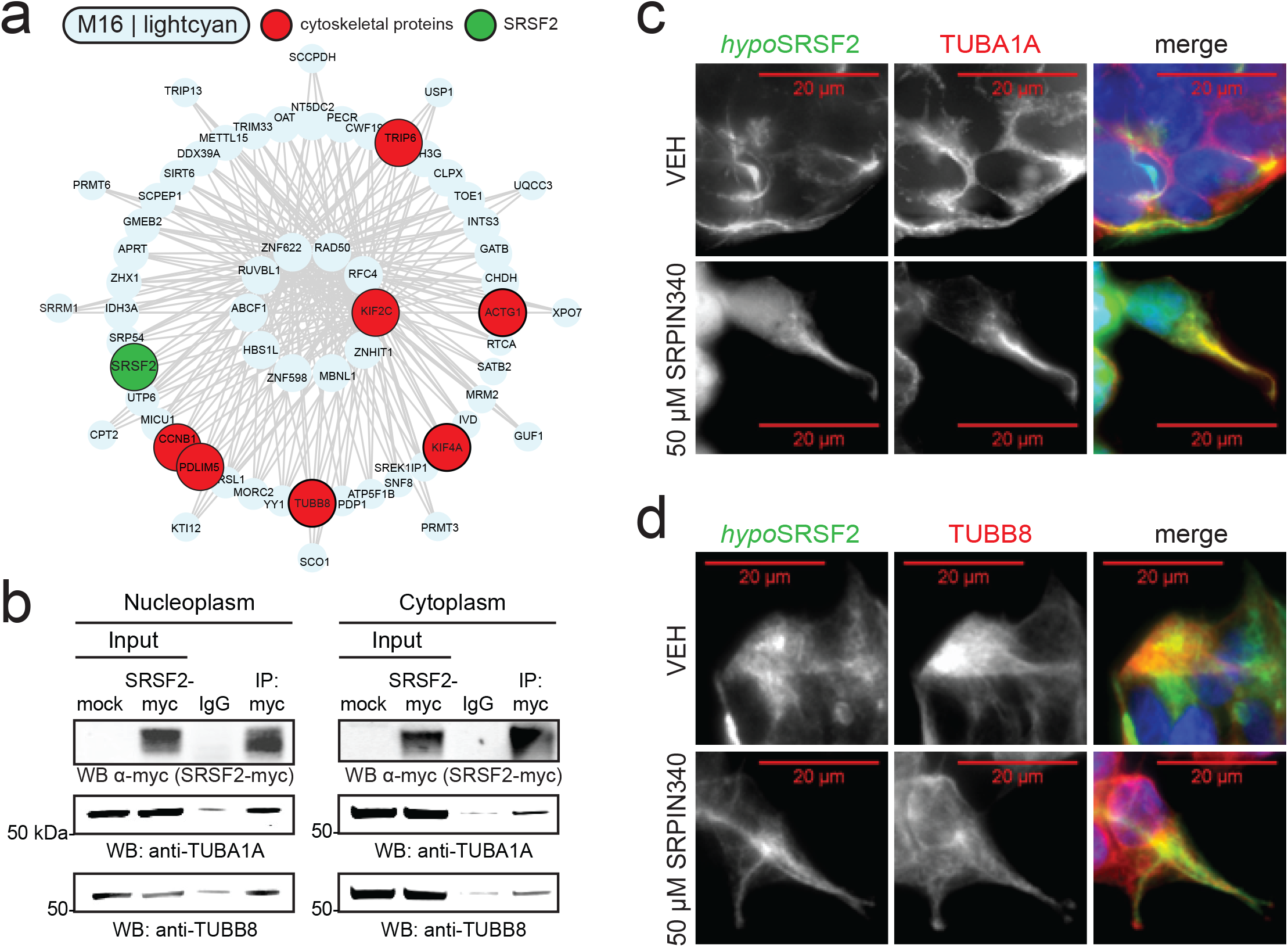
Association of SRSF2 with microtubule proteins. **(a)** I-graph of module 16 (M16) representing hub proteins and corresponding gene symbols as nodes. Node size and edges (gray) are reflective of the degree of intramodular connectivity in WGCNA (kME). SRSF2 (*green*) and cytoskeletal-associated proteins (*red*) are highlighted. **(b)** Representative western blot of an immunoprecipitation (IP) of recombinant SRSF2-myc in cytoplasm and nucleoplasm extracts isolated from HEK293 cells (*n*=3 replicates per fraction). Co-IP complexes were blotted for α-tubulin (TUBA1A) and β-tubulin (TUBB8). **(c-d)** ICC staining of HEK293 cells for hypophosphorylated SRSF2 (hypoSRSF2) (*green*) and both TUBA1A (*red*; **c**) and TUBB8 (*red*; **d**) in vehicle-cells or 50 µM SRPIN340-treated cells.

## DISCUSSION

Here we used a mass spectrometry-based proteomics approach to investigate how phosphorylation affects protein solubility and oligomerization. Using systems biology analyses we identified modules of proteins with shared biology and sequence homology that co-aggregate following dephosphorylation. Arginine-rich proteins, including SRSF2, were among the proteins with the most decreased solubility following dephosphorylation. SRSF2 was used as a paradigm to investigate the relationship between phosphorylation and protein structure and solubility. We discovered that dephosphorylation regulates higher order multimer formation of SRSF2, whereas phosphorylated SRSF2 exists largely as a monomer species *in vitro*. Inhibition of SR protein kinases within mammalian cells decreased SR protein phosphorylation, resulting in increased cytoplasmic SRSF2 granule formation, as well as cytoplasmic tubule SRSF2 structures which co-localize with cytoskeletal proteins.

SRSF2 is a well-studied classical SR protein with integral roles in constitutive and alternative splicing (30). This study highlights numerous behaviors of SRSF2 that are associated with RBP dysregulation in neurodegenerative disease, including decreased solubility, high molecular weight oligomer formation and cytoplasmic granule and tubule-like morphologies. Although mutations in SRSF2 are frequently observed in individuals with myelodysplastic syndromes (MDS) or chronic myelomonocytic leukemia (CMML) (73-77), SRSF2 has not been commonly associated with human neurodegenerative disease. Recently, however, the McKnight group demonstrated that SRSF2 does indeed form condensates *in vitro*, a hallmark feature of RBPs that aggregate in neurodegenerative disease, which importantly was reversible by phosphorylation (50,78).

Collectively, these data suggest that phosphorylation tunes SRSF2 net charge, solubility and structure by virtue of multimer disassembly *in vitro* and *in vivo* (**Fig.7**). While it is generally considered that phosphorylation promotes protein aggregation, a notable finding of this study is that arginine-rich RBPs, such as the SR and LUC7L protein families, exhibit remarkably similar aggregation patterns when dephosphorylated. Indeed, our group and others have observed that arginine-rich splicing proteins mislocalize and aggregate in AD brain (14,15,79,80). As kinases are dysregulated in AD (25), SRSF2 is a protein susceptible to solubility dysregulation in disease. Remarkably, SRSF2 was identified as a novel protein that associates with phosphorylated tau in AD brain by proteomic examination of microdissected neurofibrillary tangles (81). Our group has not identified SRSF2 as a protein with significantly increased insolubility in AD (82). However, our studies have been limited to sarkosyl-insoluble fractions only. Future studies that include internal comparisons of insoluble to soluble fractions may implicate SRSF2 and similar RBPs that are depleted from AD brain homogenate soluble fractions, and may reveal novel proteins that undergo altered solubility in AD.

**Figure 7.**
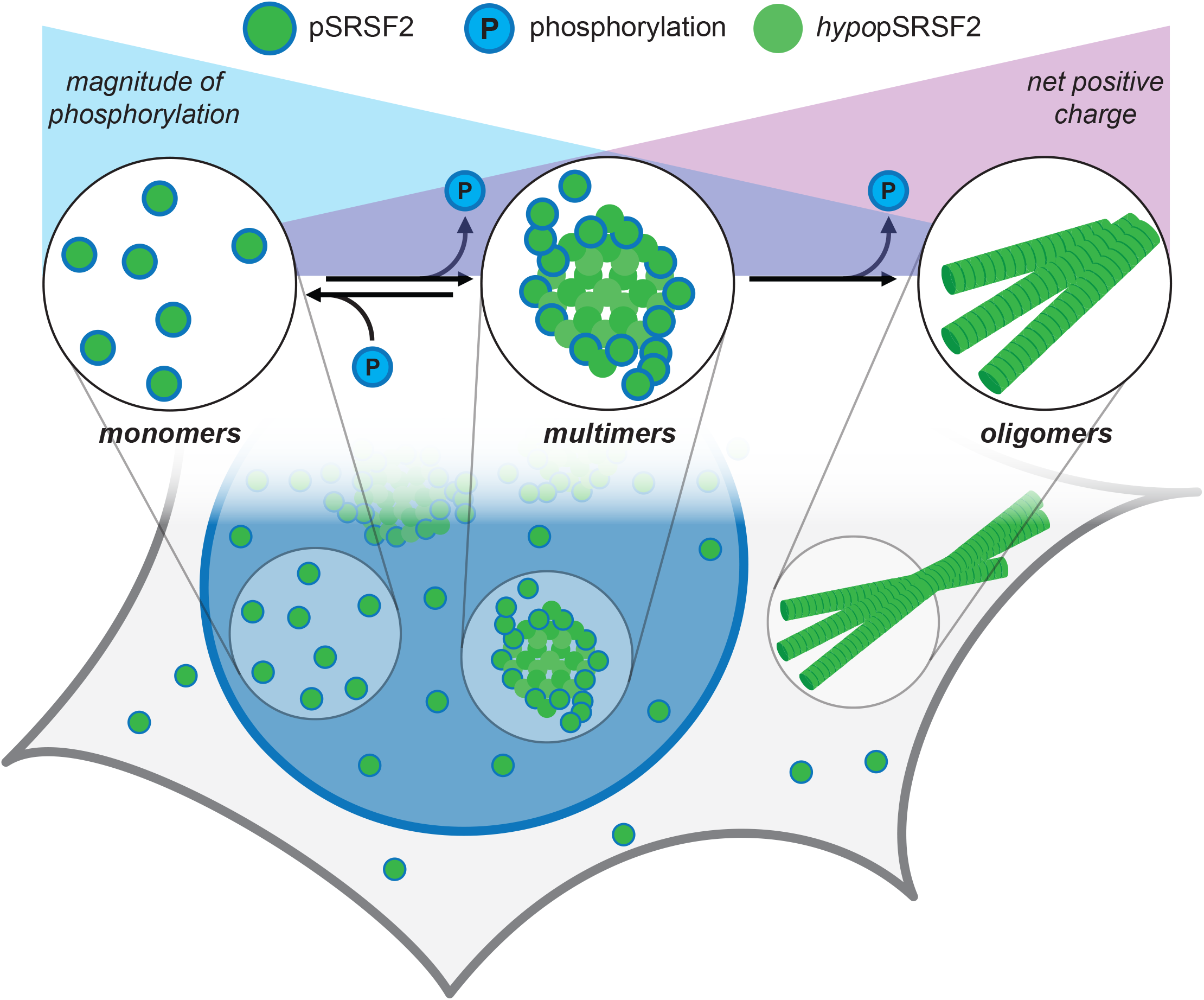
Proposed model of SRSF2 solubility and oligomerization changes regulated by phosphorylation. Equilibrium of structural states (monomer, multimer, oligomer) of SRSF2 (*green)* in varying states of phosphorylation (*dark turquoise border*). The magnitude of positive (*purple triangle*) SRSF2 net charge is illustrated, inverse to the degree of phosphorylation (*turquoise triangle*). Phosphorylation regulates higher order SRSF2 solubility, oligomerization and structure formation.

We highlight phosphorylation as a critical feature of RBPs that strongly influences protein solubility. An important consideration is the influence that similarly negatively charged RNA molecules may hold over the solubility of nuclear proteins. Future investigations that explore the role of RNA in regulating the solubility of RBPs may yield a parallel interpretation of RBP stability. Further still, studies that investigate the role of phosphorylation in the regulation of arginine-rich RBP aggregation may hold promise to reveal the mechanism underlying RBP aggregation in neurodegenerative diseases.

## EXPERIMENTAL PROCEDURES

### Materials

The primary antibodies used in this study include: rabbit polyclonal anti-myc (Cell Signaling Technology 2272S), rabbit polyclonal anti-SRSF2 (Abcam ab229473), mouse monoclonal anti-SRSF2 (Clone 1Sc-4F11, Millipore Sigma 04-1550), mouse monoclonal anti-pSRSF2 (Abcam ab11826), rabbit polyclonal anti-LUC7L (ThermoFisher 17085-1-AP), rabbit polyclonal anti-LUC7L3 (ThermoFisher PA5-53816), rat monoclonal anti-SNRPC (Abcam ab122901), rabbit polyclonal anti-SRSF1 (Abcam ab38017), rabbit polyclonal anti-TDP-43 (ProteinTech Group 10782-2-AP), rabbit anti-pS409-410 TDP-43 (CosmoBio, CAC-TIP-PTD-P02), rabbit polyclonal anti-hnRNPAB (Santa Cruz sc-98723), rabbit polyclonal anti-ZC3H18 (ThermoFisher PA5-59322), rabbit polyclonal anti-SRRM1 (ab221061), rabbit polyclonal anti-Histone H3 (Abcam ab1791), rat monoclonal anti-alpha-tubulin clone YL1/2 (MAB1864), rabbit polyclonal anti-TUBB8 (ab97880) and mouse IgM anti-pSR proteins (Clone mAb104, ATCC CRL-2067, see below isolation method). Each antibody save for mAb104 was used at a 1:1,000 dilution in blocking buffer for western blotting. Each antibody used for ICC was diluted 1:500 in normal horse serum (NHS). The pcDNA3.1-SC35-cMyc SRSF2 plasmid was a gift from Kathleen Scotto (83) (Addgene plasmid #44721).

### mAb104 Isolation

Immortalized mouse hybridomas were purchased from ATCC (CRL-2067). Cells were thawed and then passaged twice before collection. Forty-eight hours post-passaging, the cell suspension media was aliquoted into a 15 ml conical tube then spun at 250xg for 5 min. The supernatant was aspirated and cells were resuspended in 2%FBS in DMEM. Cells were cultured for 3 more days then the cell suspension was aliquoted into a 15 mL conical tube then spun at 1,500xg for 15 min. The supernatant was transferred to a fresh 15 mL conical tube, then aliquoted into microcentrifuge tubes. The supernatant aliquot volumes were then reduced under SpeedVac (Labconco 731022) to 50% of original volume and restored to the original volume with 100% glycerol. Tubes were then frozen at -20C. The antibody was diluted 1:5 in blocking buffer for western blotting.

### Local Charge Density and NCPR

Residue charge at physiological pH (7.4) was calculated using a simple algorithm modeled from EMBOSS. The residues D/E were counted with charge = -1, residues K/R with charge = +1, and H with charge = +0.5. Residues S/T/Y were counted as either charge = 0 or -2, to mimic the charge assumed after phosphorylation post-translational modification at physiological pH (63). To generate the local charge density, the average charge over a window of 21 residues starting at the C-terminus was calculated, sliding towards the N-terminus of the protein. To calculate the Net Charge per Residue (NCPR) of full proteins, the residue charges, as calculated above, were summed and this number was divided by the number of residues in the protein. The NCPR of the average protein of the known human proteome is ∼ +0.013, spanning -0.40 to 0.32 (minimum protein size = 100 residues).

### Cell Culture and Transfection

HEK293 cells were cultured in Dulbecco’s Modified Eagle Medium [DMEM, high glucose (Gibco)] supplemented with 10% (v/v) fetal bovine serum (Gibco) and 1% penicillin-streptomycin (Gibco), and maintained at 37°C under a humidified atmosphere of 5% (v/v) CO2 in air. Cells were grown to 70-80% confluency in 10 cm^2^ culture dishes and transfected with 10 μg SRSF2-myc plasmid and 30 μg linear polyethylenimine (PEI). Cells were harvested and fractionated to enrich for nucleoplasm as described below. For western blotting SRPIN340-inhibitor treated cells, HEK293 cells were incubated for 12 hours (84) with equivolume amounts of either DMSO (vehicle) or SRPIN340 (final concentration = 50 µM) starting 24 hours post-passage. Afterwards, cells were harvested in IP lysis buffer (50 mM HEPES pH7.4, 150 mM NaCl, 5% Glycerol, 1 mM EDTA, 0.5 (v/v) NP-40, 0.5% (v/v) CHAPS) and sonicated at 25% amplitude for 3 × 10 sec on/off cycles. Lysates were then cleared after a 15,600xg centrifugation step at 4°C. Supernatants were transferred to new tubes and run by SDS-PAGE followed by western blotting.

### Nucleoplasm Enrichment

This cellular extraction procedure was adapted from the Gozani group (85) and modified to include NP-40 detergent. In short, forty eight hours post-transfection, HEK293 cells were rinsed with cold PBS then scraped in PBS+1xHALT protease/phosphatase inhibitor (ThermoFisher). The cell slurry was centrifuged for 5 min at 1,000xg at 4°C to pellet cells. The supernatant was aspirated and the cells were washed with 1 mL PBS+1xHALT (100 µL taken as total fraction) and centrifuged again to pellet cells. The supernatant was aspirated and the cells were swelled in 150 µL Hypotonic Lysis Buffer+1xHALT (10 mM HEPES pH 7.9, 20 mM KCl, 0.1 mM EDTA, mM dithiothreitol (DTT), 5% Glycerol, 0.5 mM PMSF, 10 ug/mL Aprotinin, 10 ug/mL Leupeptin, 0.1% NP-40) and incubated on ice for 5 min. The sample was then centrifuged for 10 min at 15,600xg at 4°C. The supernatant was collected as the cytoplasmic fraction, and the resulting pellet was incubated in 100 µL High Salt Buffer+1xHALT (20 mM HEPES pH 7.9, 0.4 M NaCl, 1 mM EDTA, 1 mM EGTA, 1 mM DTT, 0.5 mM PMSF, 10 ug/mL Aprotinin, 10 ug/mL Leupeptin) for 30 min on ice to extract nuclei. The samples were then sonicated for 5 sec at 25% amplitude and centrifuged at 18,213xg for 10 min at 4°C. The supernatant was collected as the nucleoplasm fraction, and the pellet (chromatin fraction) was resuspended and sonicated in Nuclei lysis buffer (50 mM Tris-HCl pH 8.0, 10 mM EDTA, 1% SDS). All fractions were frozen at -70°C, and only the cytoplasm and nucleoplasm fraction were used for following applications.

### Calf Intestinal Phosphatase (CIP) Treatment

For dephosphorylation assays, nucleoplasm fractions (100 µg) were incubated with either mock (distilled water) or 50 Units of calf intestinal alkaline phosphatase (QuickCIP, NEB M0525L) for 1 hour at 37°C. Following this, Laemmli sample buffer (8% glycerol, 2% SDS, 50mM Tris pH 6.8, 3.25% beta-mercaptoethanol) was added to each sample to 1X and boiled at 95°C for 10 minutes and run by SDS PAGE. For sedimentation assays, nucleoplasm fractions (100 µg) were incubated with either distilled water (mock) or 50 Units of calf intestinal alkaline phosphatase (QuickCIP, NEB M0525L) and brought up to 100 uL with PBS. The samples were heated at 37°C for 1 hour.

### Sedimentation Assay

Following QuickCIP dephosphorylation, 50 µL of the 100 µL sample was added to polycarbonate ultracentrifuge tubes. Samples were spun at 100,000xg for 1 hour at 4°C. The supernatant (vol=50 µL) was transferred to a LoBind tube (Eppendorf 0030108442) and the insoluble pellet was resuspended in 50 µL 8M Urea and transferred to a separate LoBind tube. The pellet sample was sonicated for 1 sec on/off cycles at 25% amplitude until the pellet disappeared. For western blotting, 7.5 µL of total (pre-spin) fractions were added and 15 µL of soluble and pellet fractions were added. For mass spectrometry analysis 30 µL of soluble and pellet fractions were used.

### Western Blotting

Western Blotting was performed according to standard protocol as previously described (28). In short, samples were boiled in Laemmli sample buffer (8% glycerol, 2% SDS, 50mM Tris pH 6.8, 3.25% beta-mercaptoethanol) for 10 minutes, then resolved on a Bolt® 4-12% Bis-tris gel (Invitrogen NW04120BOX) by SDS-PAGE and semi-dry transferred to a nitrocellulose membrane with the iBlot2 system (ThermoFisher IB21001). Membranes were blocked with TBS Starting Block Blocking Buffer (ThermoFisher 37542) and probed with primary antibodies (1:1,000 dilutions) overnight at 4°C. Membranes were then incubated with secondary antibodies conjugated to either Alexa Fluor 680 or 800 (Invitrogen) fluorophores for one hour at RT. Membranes were imaged using an Odyssey Infrared Imaging System (Li-Cor Biosciences) and band intensities were calculated using Odyssey imaging software.

### Sample preparation for mass spectrometry analyses

Thirty microliters of soluble and pellet fractions were normalized to 50 µL with 8M urea buffer. Next, 10 mM Dithiothreitol (DTT) in 50 mM ammonium bicarbonate (ABC) was added to a final concentration of 1 mM and incubated for 30 min at room temperature (RT), then 50 mM iodoacetamide (IAA) in 50 mM ABC was added to a final concentration of 5 mM and incubated in the dark for 30 min at RT. Samples were then digested overnight with 1 µg of Lys-C (Wako 121-05063). Following Lys-C digestion, samples were diluted to 1M urea and digested overnight with 1 µg of Trypsin (ThermoFisher 90057). The next day samples were incubated with acidifying buffer [10% Formic Acid (FA), 1% trifluoroacetic acid (TFA)] and centrifuged for 2 min. Sample pH was verified as less than 3 using pH strips. Samples were desalted on an Oasis® PRIME HLB 10 mg plate (Oasis 186008053) and washes were flowed through columns by a 96-well Positive Pressure processor (Waters 186006961). Samples were washed first with methanol, then Buffer A (0.1% TFA). The digested samples were then loaded onto Oasis® PRIME HLB 10 mg plates (Oasis 186008053), washed with Buffer A twice, and then peptides were eluted with Buffer C [50% acetonitrile (ACN), 0.1% FA]. The elutant was lyophilized using a SpeedVac (Labconco 731022).

### Mass spectrometry analysis

Lyophilized peptides were resuspended in loading buffer (0.1% FA, 0.03% TFA, 1% ACN) and separated on a self-packed C18 (1.9 μm Dr. Maisch, Germany) fused silica column (20 cm × 75 μm internal diameter; New Objective, Woburn, MA) by a NanoAcquity UHPLC (Waters). Linear gradient elution was performed using Buffer A (0.1% formic acid, 0% acetonitrile) and Buffer B (0.1% formic acid, 80% acetonitrile) starting from 3% Buffer B to 40% over 100 min at a flow rate of 300 nl/min. Mass spectrometry was performed on an Orbitrap Fusion Lumos Mass Spectrometer in top speed mode. One full MS1 scan was collected followed by as many data-dependent MS/MS scans that could fit within a three second cycle. MS1 scans (400-1600 m/z range, 400,000 AGC, 50 ms maximum ion time) were collected in the Orbitrap at a resolution of 60,000 in profile mode with FAIMS CV set at -45. The MS/MS spectra (1.6 m/z isolation width, 35% collision energy, 10,000 AGC) were acquired in the ion trap. Dynamic exclusion was set to exclude previous sequenced precursor ions for 30 seconds with a mass tolerance of 10 ppm.

### Database Searching

Data files for the 16 samples were analyzed using MaxQuant v1.6.17.0 with Thermo Foundation 2.0 for RAW file reading capability. The search engine Andromeda (86) was used to build and search a concatenated target-decoy UniProt Knowledgebase (UniProtKB) containing both Swiss-Prot and TrEMBL human protein sequences (86,395 sequences, downloaded August 9, 2020) with 245 contaminant proteins as a parameter for the search (87). Methionine oxidation (+ 15.9949 Da), protein N-terminal acetylation (+ 42.0106 Da) and STY-phosphorylation (+ 79.966 Da) were included as variable modifications (up to 5 allowed per peptide); cysteine was assigned a fixed carbamidomethyl modification (+ 57.0215 Da). Fully tryptic peptides were considered with up to 2 miscleavages allowed in the search. A precursor mass tolerance of ± 20 ppm was applied prior to mass accuracy calibration and ± 4.5 ppm after internal MaxQuant calibration. The false discovery rate (FDR) for peptide spectral matches, proteins and site decoy fraction were all set to 1%. Phosphorylated peptides were not quantified. Protein group intensity was used for protein quantitation. The quantitation method did not consider reverse, contaminant, and by site only protein identifications, leaving 5,017 proteins for downstream analysis. The mass spectrometry proteomics data have been deposited to the ProteomeXchange Consortium via the PRIDE partner repository (88) with the dataset identifier PXD026894 on 6/23/2021.

### Weighted Gene Correlation Network Analysis of Nuclear Fractionation Proteome

Prior to network analysis, the nucleoplasm proteome (5,017 proteins) was culled to select for proteins with less than or equal to 50% missing values (intensity values in 8 out of 16 samples; 4,366 proteins). The intensity values were then log2-transformed. Missing protein intensity values were imputed using random numbers drawn from a normal distribution of observed protein intensity values of each sample (width = 0.3, down shift = 1.8, calculated separately for each column) in Perseus v1.6.15.0. The R package Weighted Gene Correlation Network Analysis (WGCNA) v1.68 was used to cluster proteins by abundance into groups of proteins with similar solubility patterns using a dissimilarity metric for clustering distance based on 1 minus the topology overlap matrix (1-TOM), a calculation based on an adjacency matrix of correlations of all pairs of proteins in the abundance matrix supplied to WGCNA (64). The weighted protein co-expression network was built using the log2-transformed intensity values using the blockwiseModules function with the following parameters: soft threshold power beta = 20, deepSplit = 4, minimum module size = 15, merge cut height = 0.07, signed network partitioning about medioids respecting the dendrogram, and a reassignment threshold of p = 0.05. GO Elite v1.2.5 python package was used as previously published to categorize summary biological functions of individual modules (89,90). Z scores were determined by one-tailed Fisher’s exact text (Benjamini-Hochberg FDR corrected) to demonstrate overrepresentation of ontologies in the nucleoplasm proteome of each module. The filters included a cutoff for Z scores as 1.96, P value cut off of 0.01 and a minimum of 5 genes per ontology.

### Differential Solubility Analysis

Proteins either differentially soluble or differentially insoluble following dephosphorylation were calculated using a two-tailed paired t-test on the fraction insolubility values according to a previous study by our group (91):

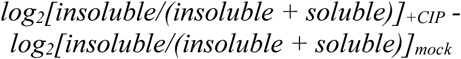

Proteins were highlighted as enriched/depleted from a fraction if greater than or equal to a two-fold change in fraction insoluble values (±log2(2)=1) was observed with a p-Value less than 0.05 (two-tailed paired t-test).

### SRPK2 in vitro kinase reaction

Human recombinant SRPK2 (0.4 ug/reaction; EMD Millipore 14-666; Glu46-end) and SRSF2 (0.8 ug/reaction; MyBioSource, Inc. MBS2029592; Thr14-end) expressed in *E. coli* were purchased and added to Kinase Buffer I (25 mM MOPS pH7.2, 12.5 mM β-glycerol phosphate, 25 mM MgCl2, 5 mM EGTA, 2 mM EDTA, 0.25 mM DTT; Abcam ab189135). Either water (mock) or ATP was added to a final concentration of 100 µM to a final volume of 25 µL and incubated for 30 min at 30°C. Reactions were then separated by denaturing or non-denaturing Blue native Gel Electrophoresis or by Isoelectric focusing (IEF).

### Blue Native Gel Electrophoresis

Blue native PAGE was carried out as described previously (16). Purified recombinant SRPK2 and SRSF2 proteins (0.4 and 0.8 µg, respectively) were incubated with blue native gel loading buffer [5% glycerol, 50mM TCEP (Sigma-Aldrich 646547), 0.02% (w/v) Coomassie G-250 (Invitrogen BN2004), 1X NativePAGE Sample Buffer (Invitrogen BN2003)] in LoBind tubes (Eppendorf 0030108442) for 30 min at room temperature. Mock and CIP-treated nucleoplasm lysate (50 µg) were incubated with the same blue native gel loading buffer and incubated on ice for 30 min. Samples and Blue Native Protein Ladder (Thermo LC0725) were loaded onto NativePAGE Bis-Tris Gels (3-12%, Invitrogen BN1001BOX) and gel electrophoresis was performed using anode NativePAGE Running Buffer (Invitrogen BN2001) and cathode buffer (Invitrogen BN2002) with additive (Invitrogen BN2004) and run for 15 min at 150V. The Dark Blue Cathode Buffer was then interchanged with Light Blue Cathode Buffer, proceeding for another 90 min at 150V. Following this, the gel was gently rocked in 50 mM Tris-HCl, pH7.5 and 1% SDS for 30 min and transferred using the semidry iBolt transfer system (Invitrogen) with PVDF membrane (Invitrogen IB24002) for 7 min at 20V. Following transfer, the PVDF membrane was rocked in 8% acetic acid solution for 5 min, rinsed with distilled water, air dried, then rinsed with methanol. The membrane was then blocked, western blotted and imaged on the Odyssey Infrared Imaging System (Li-Cor Biosciences).

### Isoelectric Focusing (IEF) Gel Electrophoresis

*In vitro* kinase reactions (10 µL; 400 ng SRSF2, 200 ng SRPK2) were mixed with IEF Sample Buffer pH 3-10 (Fisher Scientific LC5311) and dH2O. Samples were loaded onto Novex IEF pH 3-10 gels (ThermoFisher EC6655BOX) and gel electrophoresis was run with 1X IEF Anode Buffer (ThermoFisher LC5300) and 1X Cathode Buffer pH 3-100 (ThermoFisher LC5310). Protein marker ladder was included to benchmark protein pI (ThermoFisher 3921201). The gel was run by 100V constant for 1 hr, 200V constant for 1 hr, then 500V for 30 min. Following electrophoresis, the gel was fixed in 12% TCA containing 3.5% sulfosalicylic acid. After fixation, the gel was washed 3 x dH2O. The gel was then stained overnight with Coomassie G-250.

### Immunocytochemistry

Immunocytochemical staining was carried out as previously described (92). HEK293 cells were grown on Nunc Lab-Tek II Chamber Slide systems (ThermoFisher 154534PK). Twenty four hours post-passage, HEK293 cells were incubated with equivolume amounts of either DMSO (vehicle) or SRPIN340 (final concentration = 50 µM). Cells were incubated at 37°C under a humidified atmosphere of 5% (v/v) CO2 in air for 4 hours. Cell media was aspirated then washed 3 × warm sterile PBS for 5 min each. Cells were then fixed with 4% paraformaldehyde (Electron Microscopy Sciences 15713S) diluted in PBS for 45 min at RT and afterwards washed 3 × warm sterile PBS for 5 min each. Cells were then incubated with 0.05% Triton X-100 diluted in PBS for 20 min at RT. Afterwards, cells were blocked with 10% normal horse serum (NHS) in PBS for 45 min at RT. The blocking solution was aspirated and excess liquid was removed with a kimwipe. The cells were then incubated in primary antibody diluted in 2% NHS/1xPBS overnight at 4°C. The next day, cells were washed 3 × warm sterile PBS for 5 min each. Cells were then incubated in secondary antibody diluted in 2% NHS/1xPBS for 1 hour at RT then washed 3 × warm sterile PBS for 5 min each. PBS was aspirated and a droplet of DAPI-containing mounting media (Abcam ab104139) was placed on cells which were coverslipped and sealed with clear nail polish. Images were captured on a Keyence BZ-X810 widefield laser scanning microscope (Keyence). At least 14 images were taken for each condition per biological replicate at random areas of the slide.

### Imaging Quantification and Statistical Analysis

Images were analyzed with FIJI. Graphs were developed with GraphPad Prism. Over 100 cells each were counted (DAPI+ nuclei) and analyzed per three biological replicates. Then, the number of SC35+ (pSRSF2) granules were counted in each image and divided by total cells observed. The average number of granules per cell per image (*n*=14) was collected for each biological replicate. At least 120 cells were counted in each condition per replicate. The number of cytoplasmic granules per cell was averaged for each replicate and conditions were statistically compared using a two-tailed paired t-test. To measure the percent of cells with cytoplasmic SRSF2 tubules, the total number of DAPI+ nuclei were counted as the number of cells per image. A total of 15 images per biological replicate were recorded, with four biological replicates analyzed. Then, the number of cells that contained an SRSF2 tubule morphology were counted. The number of SRSF2 tubule-containing cells were then divided by the number of DAPI+ nuclei and expressed as a decimal. The number of SRSF2 tubules per cell was averaged in each replicate and the vehicle and drug conditions were statistically compared using a two-tailed paired t-test.

## ACKNOWLEDGMENTS

We thank Anita Corbett and Zachary T. McEachin for their helpful comments in discussion of this work.

## Author contributions

SK designed and carried out the experiments. DD, LP, LY and NTS advised experimental procedures. SK, ED and CH performed the data analyses. SK and ED did the computational analyses. SK drafted the manuscript and figures. SK and NS wrote and edited the manuscript. SK, ED, CH, LY, LP, DD and NTS reviewed and edited the manuscript. NTS carried out funding acquisition. All authors contributed to the article and approved the submitted version.

## FOOTNOTES

Support for this research was provided by funding from the National Institute on Aging (R01AG053960, R01AG061800, and RF1AG062181), the Accelerating Medicine Partnership for AD (U01AG046161 and U01AG061357). S.K. was supported by the Training Program in Biochemistry, Cell and Developmental Biology (T32GM008367) and Emory Neurology Department Training Fellowship (5T32NS007480-19).

## Supplemental Figures

**Supplemental Figure S1.**
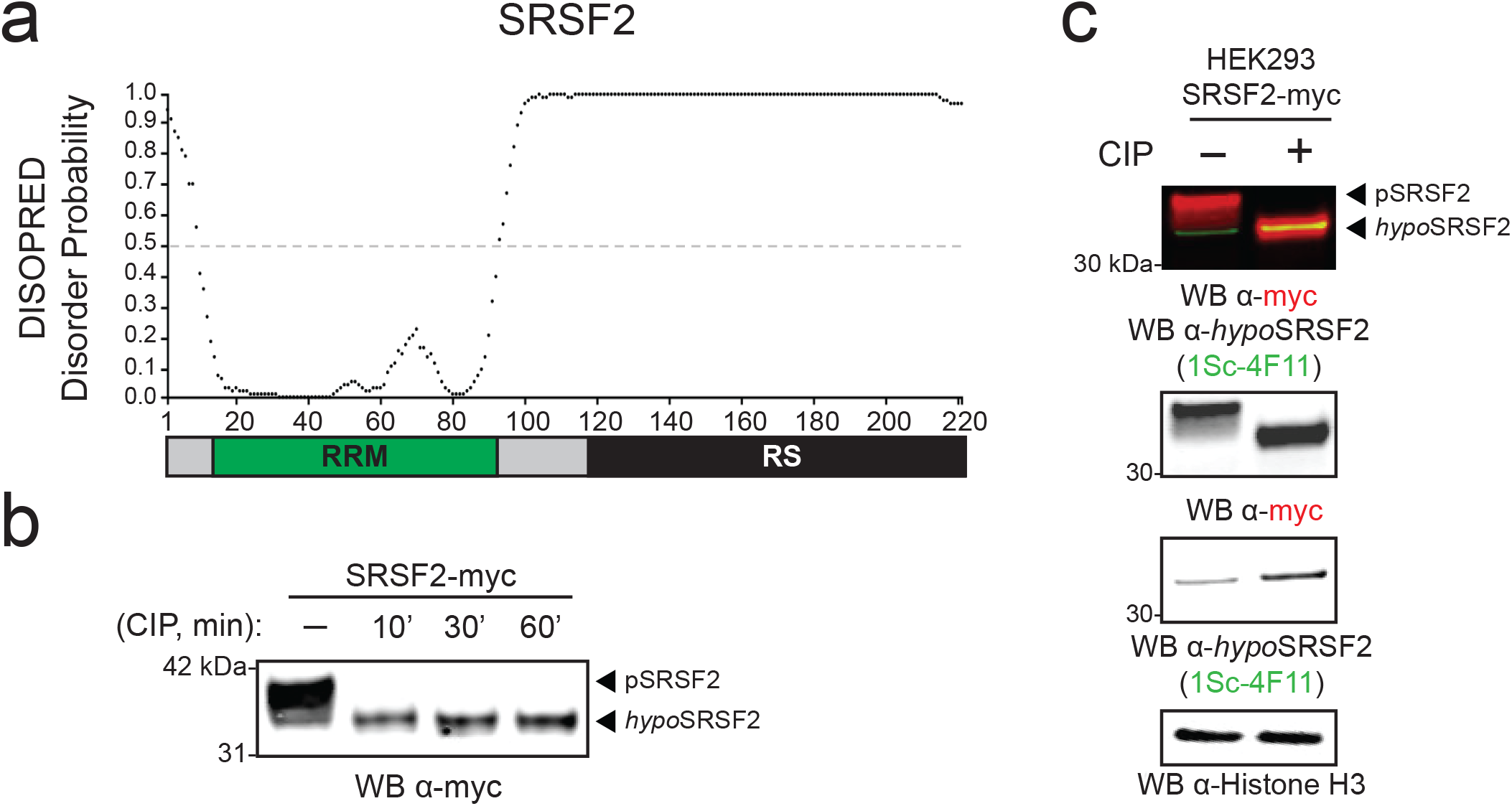
The arginine-/serine-rich (RS) domain of SRSF2 is predicted to be highly disordered. The DISOPRED 3.0 algorithm predicts intrinsically disordered regions of proteins based on primary sequence alone. The protein architecture of SRSF2 below is sized to match the x-axis of the disorder plot. SRSF2 harbors an N-terminal RNA-recognition motif (RRM) domain (*green box*) and a C-terminal arginine-/serine-rich (RS) domain (*black box*). **(b)** Nucleoplasm extracts of HEK293 cells expressing recombinant SRSF2-myc protein were incubated with either distilled calf intestinal alkaline phosphatase (+CIP) for increasing time lengths (10 min, 30 min, 1 hour) or water (-CIP) or at 37°C and separated by denaturing SDS-PAGE and immunoblotted for the myc tag. **(c)** Nucleoplasm fractions of HEK293 cells transiently-expressing SRSF2-myc were treated with either dH2O (-) or CIP (+), separated by SDS-PAGE and immunoblotted for myc tag (*red*) and hypophosphorylated SRSF2 (*green*, hypoSRSF2) as well as Histone H3 for loading control. Increased electrophoretic mobility of SRSF2-myc was observed in +CIP treatment, along with increased labeling by hypoSRSF2-specific antibody (1Sc-4F11).

**Supplemental Figure S2.**
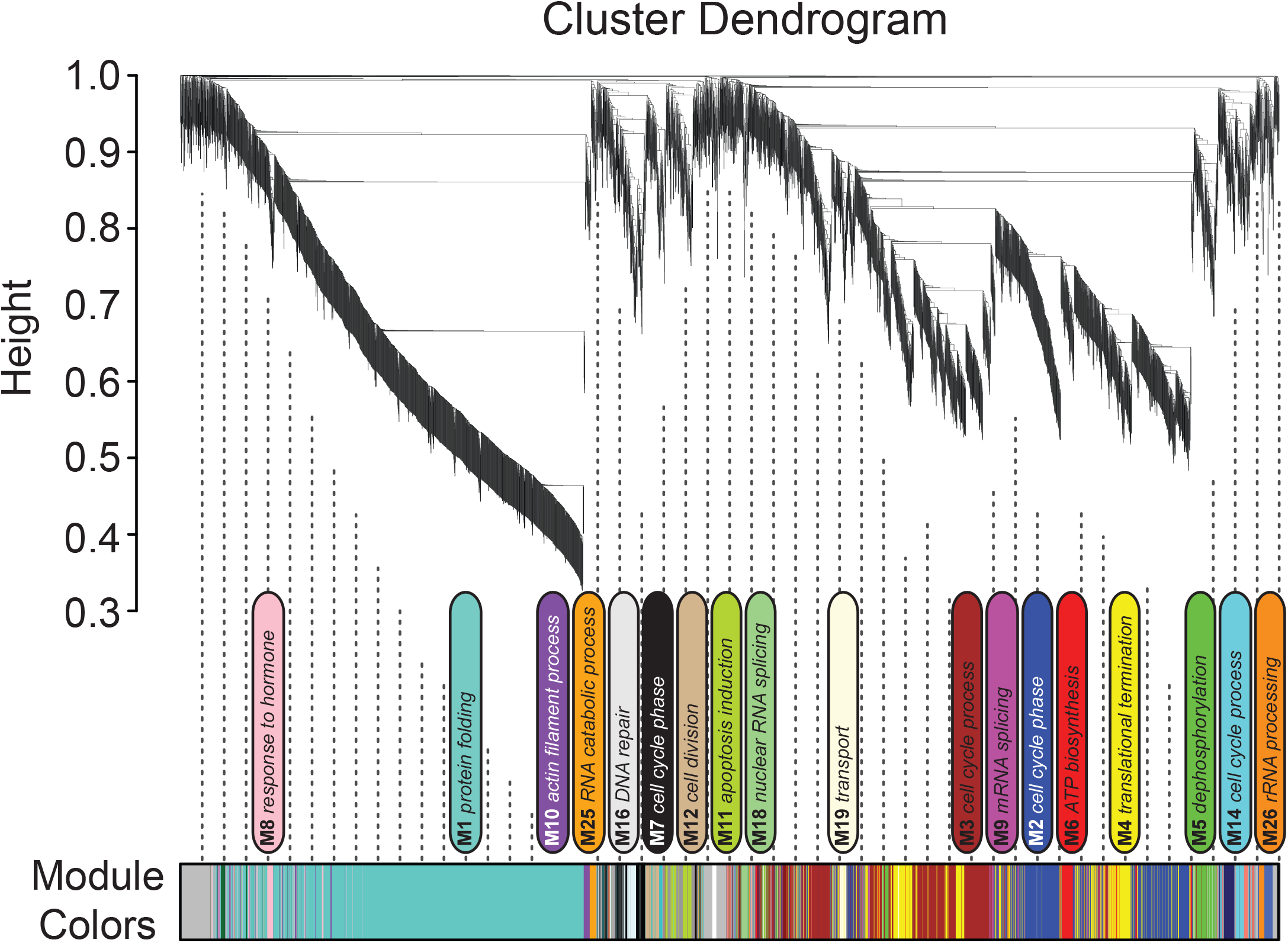
The nuclear proteome was separated into discrete groups by weighted gene correlation network analyses. Weighted Gene Correlation Network Analysis (WGCNA) cluster dendrogram groups all proteins (*n*=4,366) measured by hierarchical clustering into 27 different protein modules (M1-M27). The top generalizable biological process gene ontology term was given as a title to each module.

**Supplemental Figure S3.**
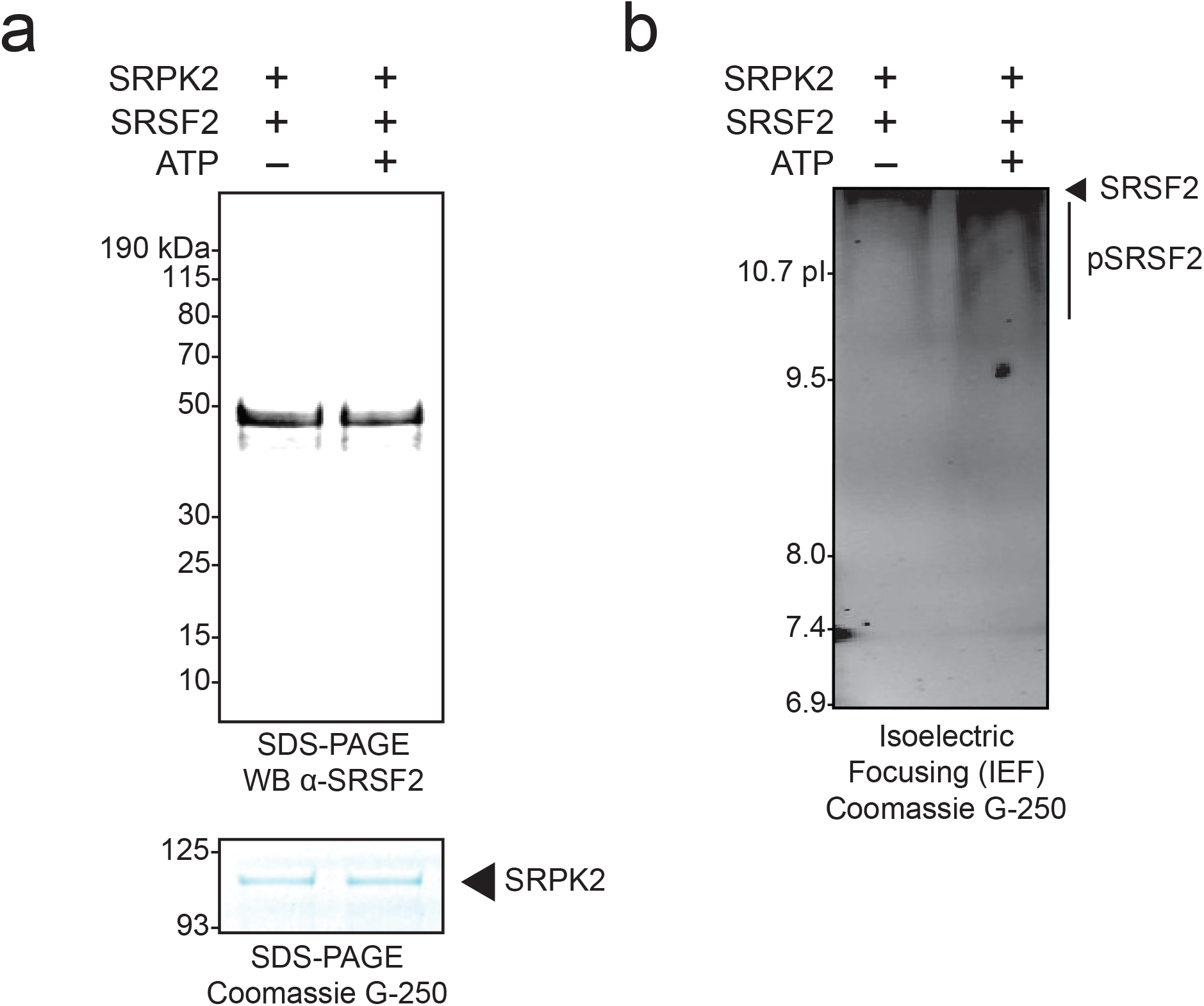
Validation of *in vitro* SRPK2-SRSF2 kinase reactions. (**a-b**) Serine-/arginine protein kinase 2 (SRPK2) and SRSF2, expressed and purified from *E. coli* were mixed in the presence or lack thereof (-/+, respectively) ATP and incubated at 30°C for 30 min and run by denaturing SDS-PAGE or isoelectric focusing (IEF) gel electrophoresis. **(a)** Equal loading of SRPK2 was validated by Coomassie staining (∼110 kDa). **(b)** The pI of non-phosphorylated SRSF2 (-ATP) is 12.4, while SRSF2 co-incubated with SRPK2 and ATP had an apparent pI range of 10-12.4.

